# Chemical genetics reveals cross-activation of plant developmental signaling by the immune peptide-receptor pathway

**DOI:** 10.1101/2024.07.29.605519

**Authors:** Arvid Herrmann, Krishna Mohan Sepuru, Hitoshi Endo, Ayami Nakagawa, Shuhei Kusano, Pengfei Bai, Asraa Ziadi, Hiroe Kato, Ayato Sato, Jun Liu, Libo Shan, Seisuke Kimura, Kenichiro Itami, Naoyuki Uchida, Shinya Hagihara, Keiko U. Torii

**Author notes:** Contributed equally as co-first authors (Arvid Herrmann, Krishna Mohan Sepuru). Current Address: MCDB, University of Michigan, Ann Arbor, MI 48109.

## Abstract

Cells sense and integrate multiple signals to coordinate development and defence. A receptor-kinase signaling pathway for plant stomatal development shares components with the immunity pathway. The mechanism ensuring their signal specificities remains unclear. Using chemical genetics, here we report the identification of a small molecule, kC9, that triggers excessive stomatal differentiation by inhibiting the canonical ERECTA receptor-kinase pathway. kC9 binds to and inhibits the downstream MAP kinase MPK6, perturbing its substrate interaction. Strikingly, activation of immune signaling by a bacterial flagellin peptide nullified kC9’s effects on stomatal development. This cross-activation of stomatal development by immune signaling depends on the immune receptor FLS2 and occurs even in the absence of kC9 if the ERECTA-family receptor population becomes suboptimal. Furthermore, proliferating stomatal-lineage cells are vulnerable to the immune signal penetration. Our findings suggest that the signal specificity between development and immunity can be ensured by MAP Kinase homeostasis reflecting the availability of upstream receptors, thereby providing a novel view on signal specificity.

## INTRODUCTION

Living organisms must sense numerous environmental and endogenous cues and integrate information to grow and survive. For effectively allocating their resources, plants balance their growth and defence. As such, how the growth and defence signal transduction pathways crosstalk while maintaining specificities remains an important question. It is well established that plants use a battery of cell-surface receptors, known as receptor kinases (RKs) to perceive signals (Shiu and Bleecker, 2001a, b). Among them, those RKs with extracellular leucine-rich repeats (LRR-RKs) mediate developmental and environmental/immunity signals and elicit the responses accordingly (DeFalco and Zipfel, 2021; Dievart and Clark, 2004; Tang et al., 2017; Torii, 2004). In the past decades, ligand binding, activation mechanisms, and downstream signal transduction pathways of LRR-RKs have been studied extensively (Chakraborty et al., 2019; DeFalco and Zipfel, 2021; Hohmann et al., 2017). Increasing efforts have been made to dissect the mechanism of signal crosstalk between development and defence, which happens at multiple levels from the segregation of receptor complexes at the plasma membrane to the regulation of gene expressions (Fontes, 2024; Ortiz-Morea et al., 2020). Still, understanding how each RK signaling pathway maintains signal specificity remains largely enigmatic.

The developmental patterning of stomata, cellular valves on the land plant epidermis for photosynthetic gas exchange and water control, is enforced by the ERECTA-family of LRR-RKs, ERECTA, ERECTA-LIKE1 (ERL1), and ERL2 (Shpak et al., 2005). These receptors perceive a family of secreted peptides, EPIDERMAL PATTERNING FACTORs (EPFs)/EPF-LIKEs (EPFLs), to synergistically enforce proper stomatal patterning (Hara et al., 2007; Hara et al., 2009; Lee et al., 2012). Upon ligand binding, the activated ERECTA-family receptors form a receptor complex with BRI1 ASSOCIATED KINASE1 (BAK1)/SOMATIC EMBRYOGENESIS RECEPTOR KINASEs (SERKs) to transduce cellular signals (Meng et al., 2015). The LRR receptor-like protein (RLP) TOO MANY MOUTHS (TMM) associates with ERECTA-family receptors to discriminate different EPF/EPFL peptides in a context-specific manner (Abrash and Bergmann, 2010; Abrash et al., 2011; Lee et al., 2015; Lin et al., 2017; Nadeau and Sack, 2002). The signal is then transduced to a MAPK cascade, comprising MAPKKKs YODA (also known as MAPKKK4), two redundant MAPKKs MKK4/MKK5, and two redundant MPK3/MPK6 as MAPKs (Bergmann et al., 2004; Lampard et al., 2008; Wang et al., 2007). The activated MPK3/6 inhibit stomatal differentiation via direct binding to a basic-helix-loop-helix (bHLH) transcription factor, SCREAM (SCRM, also known as ICE1), and subsequent phosphorylation and degradation of bHLH heterodimers of SCRM and SPEECHLESS (SPCH) (Kanaoka et al., 2008; Lampard et al., 2008; Putarjunan et al., 2019; Wang et al., 2007). The *SPCH* paralogs, *MUTE* and *FAMA*, act sequentially to promote the differentiation of stomatal precursor cells, meristemoids and then guard mother cells (Herrmann and Torii, 2021). Recent genetic studies indicate that the receptor-like cytoplasmic kinases (RLCKs) BRASSINOSTEROID SIGNALING KINASE1 and 2 (BSK1/2)(Tang et al., 2008) act downstream of ERECTA-family, likely bridging the receptor complex to the MAPK cascade (Neu et al., 2019).

Increasing evidence show that, whereas the upstream ligand-receptor pairs and downstream transcription factors of the LRR-RK pathways are unique, their intermediate signaling components are shared among the different pathways (Bender and Zipfel, 2023; Chen and Torii, 2023). The EPF-ERECTA-family pathway largely shares co-receptor and downstream MAPK components with the well-established immunity pathway mediated by the FLAGELLIN SENSING2 (FLS2) LRR-RK (Asai et al., 2002; Chinchilla et al., 2007; Meng et al., 2015; Wang et al., 2007; Zipfel et al., 2004). FLS2 perceives the bacterial flagellin peptide flg22, forms an active receptor complex with BAK1, and parallelly activates two downstream MAPK cascades: one mediated by MEKK1, MKK1/2, and MPK4 to prevent autoimmunity, and the other mediated by MAPKKK3/5, MKK4/5, and MPK3/6 that phosphorylate transcription factors (e.g. WRKY) for defence gene expression (Asai et al., 2002; Bender and Zipfel, 2023; Chinchilla et al., 2007; Gao et al., 2008; Zhang et al., 2012b; Zipfel et al., 2004). MKK4/5 and MPK3/6 are shared by both ERECTA-family and FLS2 pathways. How could each RK pathway maintain signal specificity while sharing the same components? Initially, stomatal development and immunity pathways are predicted to use distinct MAPKKKs, YODA and MAPKKK3/5, respectively (Bergmann et al., 2004; Bi et al., 2018), which may determine the signal specificity. Indeed, it was reported that ERECTA-family and FLS2 signaling pathways maintain specificity via antagonistic interactions between YODA (MAPKKK4) and MAPKKK3/5 (Sun et al., 2018). However, two more recent studies show that YODA (MAPKKK4), MAPKKK3, and MAPKKK5 possess overlapping functions in stomatal development and immunity, thereby arguing against their mutually inhibitory roles (Liu et al., 2022; Wang et al., 2022). These contrasting observations may stem from complex genetic redundancy and pleiotropy/lethality in these mutants, which often hamper the dissection of intricate signaling pathways.

Chemical genetics offers an unbiased approach to investigate signal transduction pathways in cells to whole organisms through perturbations of biological targets (e.g. proteins) by small molecules (Hicks and Raikhel, 2009; McCourt and Desveaux, 2010; Spring, 2005). The controllable nature of chemical application, both in dose and timing, may enable disentanglement of signaling pathways. With this in mind, we performed a large-scale chemical genetics screen and identified a small compound, kC9 (HNMPA), as a potent inducer of stomatal development via interfering with the EPF-ERECTA signaling pathway. Further mechanistic studies using available genetic mutant resources, chemical derivatization and structure-activity relationship analyses, as well as biochemical and biophysical binding assays backed up with docking modeling show that kC9 is a novel inhibitor of MAPK, MPK6. Intriguingly, we discovered that kC9’s effects in increasing stomatal development is fully nullified by the activation of the flg22-FLS2 signaling pathway. This surprising signal cross-activation of stomatal development by the immune signaling pathway occurs even in the absence of kC9, if functional population of ERECTA-family receptors becomes limited. Detailed time-course studies further uncovered the developmental window whereby stomatal-lineage cells are most vulnerable to the flg22-FLS2 immune signal activation. Our findings suggest that the signal specificity between two distinct signaling pathways in development and immunity can be ensured when they are in a fully activable state, thus challenging the conventional view of signal specificity.

## RESULTS

### A small molecule promotes stomatal development via interfering with a canonical signaling pathway

To identify novel chemical compounds with profound effects on stomatal development, we performed a phenotype-based screen utilizing our pipeline of curated small molecules on Arabidopsis seedlings expressing guard-cell-specific GFP marker (Fig. 1A; See Methods). Among ∼10,000 compounds screened, we identified hydroxy-2-napthalenymethylphosphonicacid (HNMPA, from now referred to as kC9), as a potent inducer of stomatal differentiation, resulting in severe stomatal clustering (Fig. 1; Fig. S1). HNMPA is a known inhibitor of the human insulin receptor with weak/modest efficacy (Baltensperger et al., 1992). However, its action or targets in plants are unknown. Consistent with increase in the number of stomata (Fig. S1D, E), number of cells expressing stomatal-lineage markers *TMMpro::GUS-GFP* (Figure 1B) and *MUTEpro::nucYFP* (Fig. 1C) are substantially increased. The effect of kC9 is dose-dependent, significantly increasing the number of stomata as its concentration increases (Fig. 1C-E).

**Figure 1.**
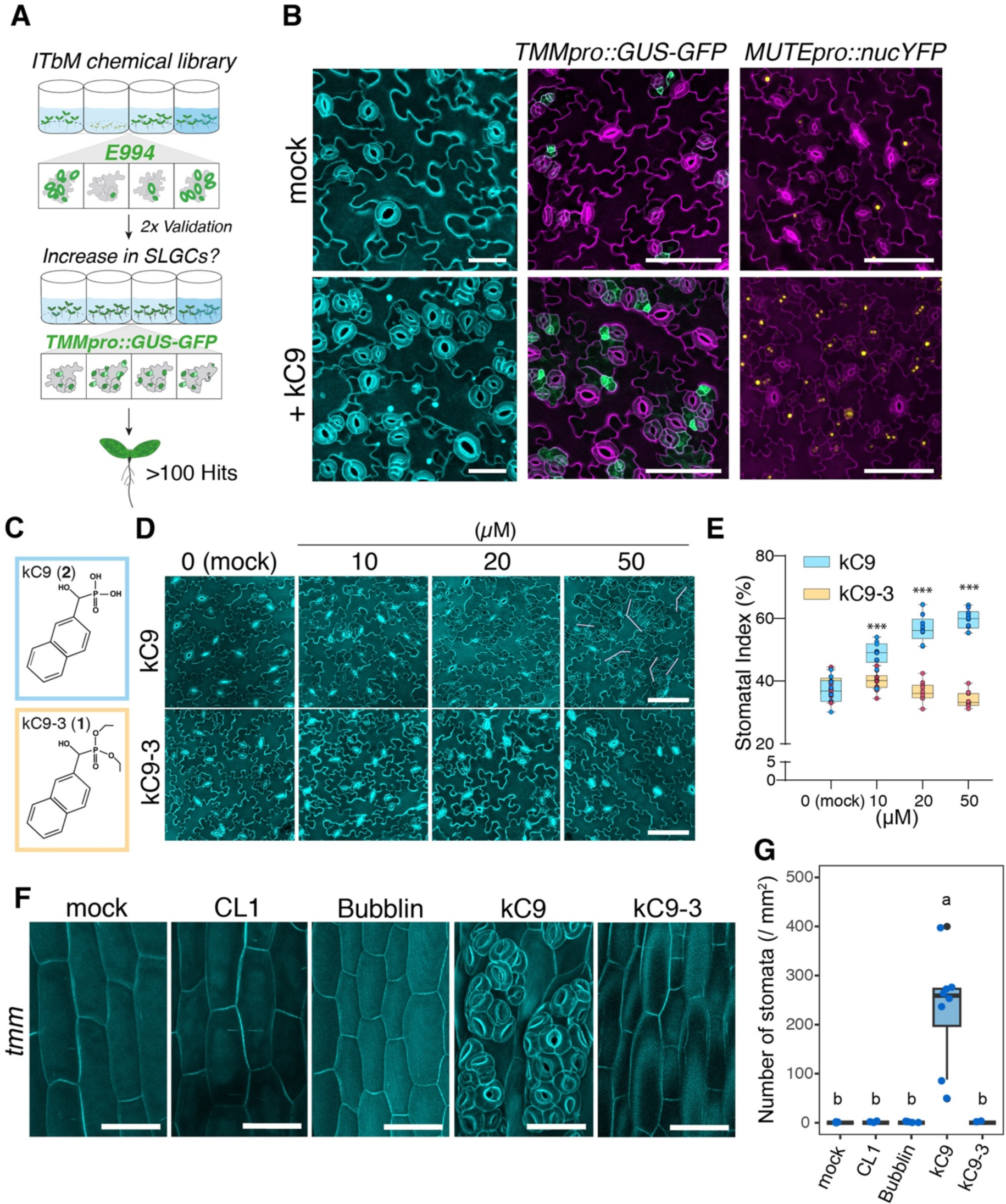
kC9 increases stomata formation via impacting the cell-cell signalling pathways. (A) Schematics of a phenotype-based screen using a synthetic compound library. Seedlings expressing the guard cell marker *E994* were screened for increased number of stomata. The phenotype was then validated by two different stomatal marker lines (*E994* and stomatal-lineage marker, *TMMpro::GUS-GFP*). (B) Effects of kC9 on stomatal differentiation. Shown are representative images of cotyledon abaxial epidermis from 9-day-old wild type (left), *TMMpro::GUS-GFP* (middle), and *MUTEpro::nucYFP* (right) seedlings treated with mock (top) or 50 µM kC9. Note the vast increase of fluorescent markers upon kC9 treatment. Cell peripheries are visualized with PI. Scale bars = 40 µm (left) and 100 µm (middle and right). (C) Chemical structure of kC9 (top; compound 2) and its non-bioactive analogue kC9-3 (bottom, compound 1). (D) Representative images of cotyledon abaxial seedling epidermis treated with kC9 or kC9-3 at 0 (mock), 10, 20, or 50 µM. Stomatal clusters are observed at 50 µM (pink brackets). Scale bars, 100 µm. (E) Quantitative analysis of stomatal index of kC9 or kC9-3 treatment in (F). n=10. Student’s unpaired t-Test, with a two-tailed distribution, kC9 vs kC9-3 per concentration, mock: NS p = 0.4621529; 10 µM:***p = 0.000298551, 20 µM: ***p = 1.2829E-09; 50 µM: *** = 7.292E-14. (F) kC9 reverts the absence of stomata on *tmm* hypocotyls. Shown are representative images of hypocotyl epidermis of 9-day-old *tmm-KO* mutant plants treated with either mock, 25µM CL1, 10 µM Bubblin, 50 µM kC9, or 50 µM kC9-3. Scale bar = 50 µm. kC9 treatment counteracts the absence of stomata in *tmm* and triggers severe stomatal clusters. (G) Quantitative analyses of stomatal density per 1 mm^2^ of hypocotyl epidermis from the *tmm* seedlings grown in the presence of mock or treated with chemical compounds as indicated in (F). One-way ANOVA followed by Tukey’s HSD analyses were performed. Letter (a or b) indicate groups that are statistically different. n = 8

To characterize the structure-activity relationship (SAR) of kC9, we subsequently synthesized a series of kC9 analogues and tested them for their bioactivity (Fig. 1C-E, Fig. S1 D, E, Document S1). kC9 (compound **2**) consists of two aryl rings (naphthalene) with a phosphonic acid through a hydroxy-methyl group (Fig. 1C, Document S1). Through chemical synthesis, we replaced the side chains (kC9-3 (**1**), kC9-A (**10**), kC9-11 (**11**), and kC9-13 (**13**)) or added an additional naphthalene to its predecessor (kC9-8 (**6**)) (Fig. S1. See Document S1 for details of synthesis and NMR analyses). While kC9 can increase the SI in a dose-dependent manner, kC9-3, in which all hydroxyl chains in the phosphoric acid moiety are replaced by methoxyl groups, failed to increase the number of stomata. Likewise, kC9-8 did not increase the number of stomata (Fig. S1 D-EG) whereas the remaining kC9 analogs exhibit bioactivity. Taken together, these results suggest that the size of aryl rings (kC9-8) and the phosphonic acid (kC9-3) are crucial for the bioactivity of kC9 to promote stomatal development.

We sought to test whether kC9 triggers stomatal development via impacting the canonical EPF-ERECTA family signaling pathway. It is known that whereas *tmm* mutant cotyledon and leaves of form stomatal clusters like *erecta (er) erl1 erl2* triple mutant, *tmm* hypocotyl is devoid of stomata (Geisler et al., 1998; Shpak et al., 2005). This is due to the role of TMM in safeguarding the inappropriate activation of ERECTA-family RKs (Abrash and Bergmann, 2010; Abrash et al., 2011; Torii, 2012; Uchida et al., 2012). Genetically, loss-of-functions in *ERECTA*-family or *EPF/EPFL* genes can ‘bring back’ stomatal differentiation on *tmm* hypocotyl epidermis (Abrash and Bergmann, 2010; Abrash et al., 2011; Shpak et al., 2005). We therefore predicted that if kC9 acts on the EPF/EPFL-ERECTA signaling pathway, its application can revert the *tmm* hypocotyl phenotype. Intriguingly, kC9 application significantly increases the stomatal development, conferring severe stomatal clustering on stomata-forming cell files whereas its inactive analog kC9-3 has no effects (Fig. 1F, G). Unlike kC9, neither the previously reported small molecule CL1, which increases stomata (Ziadi et al., 2017) nor Bubblin, which triggers asymmetric division defects and stomatal cluster formation (Sakai et al., 2017), can resume stomatal differentiation on *tmm* hypocotyls (Fig. 1F, G). Therefore, we conclude that kC9 must act within the EPF/EPFL-ERECTA-family signaling pathways.

### kC9 targets stomatal MAPK cascade

Our analyses of *tmm* mutant hypocotyls uncovered that kC9 potentially acts within the canonical stomatal cell-cell signaling pathway (Fig. 1F, G). To locate the exact target of kC9, we employed a chemical-genetic approach and treated different stomatal pathway mutants with kC9 (Fig. 2, Fig. S2). Single or higher-order mutants impaired in the peptide-receptor-signaling complex *epf1 epf2*, *er erl1 erl2, tmm* as well as *serk1 serk2 bak1* triple mutant all display a significant increase in the stomatal index upon kC9 application compared to their respective mock controls (Figs. 2B, 2C, S2A, S2B). Among the stomatal transcription factor mutants, kC9 treatment had no effects on *spch* mutant epidermis (Fig. S2C). Whereas kC9 vastly increased the arrested meristemoids and GMC tumours in *mute* and *fama,* respectively, no stomatal differentiation occurred (Fig. S2C), indicating that kC9 cannot promote stomatal differentiation in the absence of *SPCH/MUTE/FAMA*. Combined, the results suggest that kC9 affects downstream of the peptide-receptor signaling module but upstream of the bHLH master TF. As reported previously (Lee et al., 2012), induced overexpression of *EPF1* (*Est::EPF1*) or *EPF2* (*Est::EPF2*) severely inhibits stomatal development. Notably, the kC9 application has minimal effects on the inhibition of stomatal development when *EPF1/2* are overexpressed (Fig.2 D, E), indicating that activated EPF-ERECTA-family signaling can counteract the effects of kC9.

**Figure 2.**
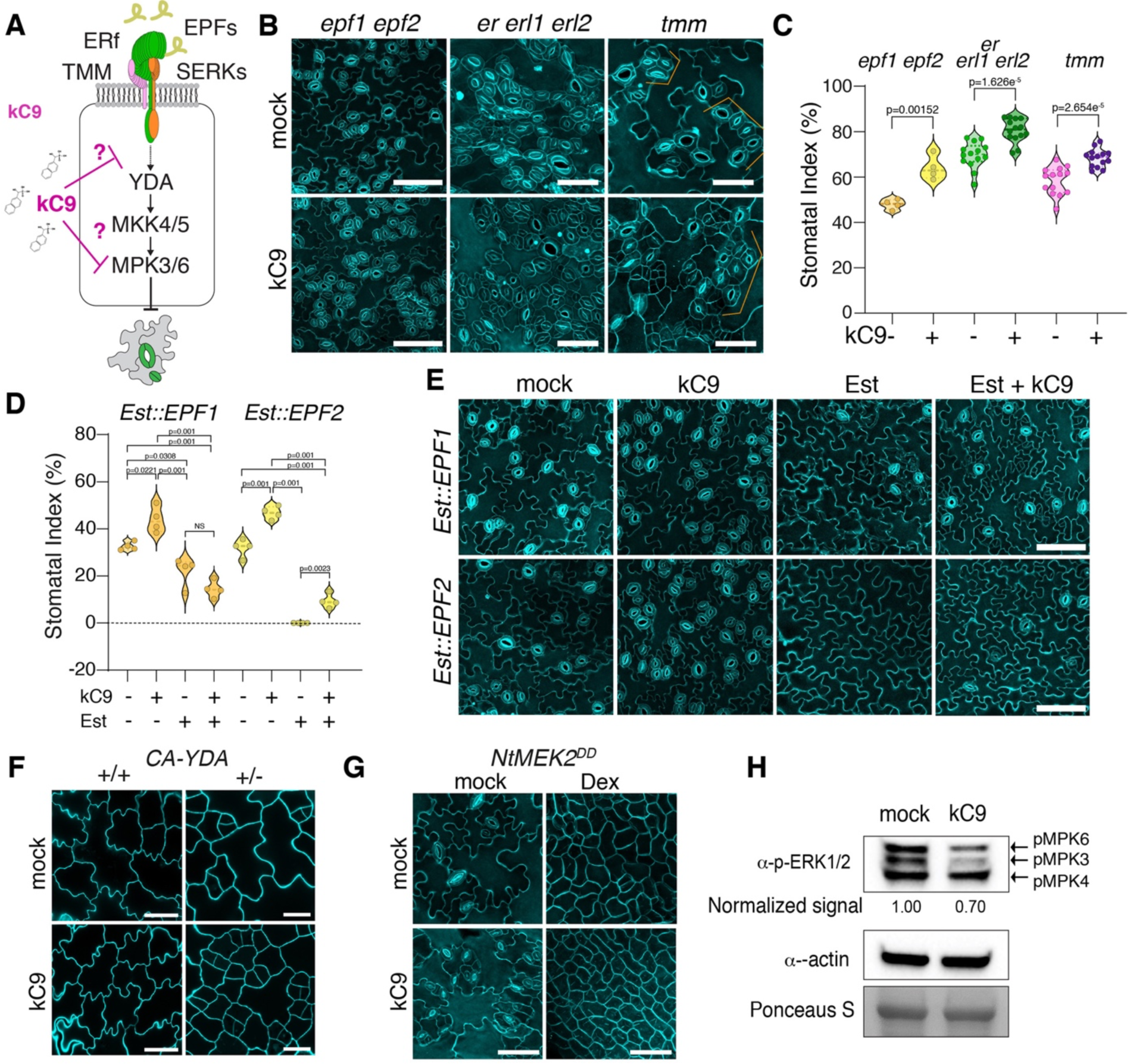
kC9 acts at the point of the MAPK. (A) Cartoon of kC9 action (red T-bar) on the stomata peptide-receptor signalling pathways. Arrow, activation. (B) Representative confocal images of the seven-day-old abaxial cotyledon epidermis from *epf1 epf2* (left), *er erl1 erl2* (middle), and *tmm* (right) seedlings either treated with either mock (top) or 50 µM kC9. Scale bars = 50 µm. (C) Stomatal Index of *epf1 epf2* (n = 4, **p = 0.0015) *er105 erl1-2 erl2-1* (n = 14, ****p =1.6256E-05) and *tmm* (n = 14, ****p = 2.6540E-05) seedlings either treated with either mock (top) or 50 µM kC9. Two-tailed unpaired Student’s t-test was performed. (D-E) Induced overexpression of either EPF1 or EPF2 counteracts the activity of kC9 on stomatal development. (D) Stomatal index of the seven-day-old abaxial epidermis from *Est::EPF1* and *Est::EPF2* seedlings treated with either mock, 50 µM kC9, 5 µM oestradiol (Est) or both. Note that after the induction of EPF1 or EPF2, the stomatal index is significantly reduced (n = 4). One-way ANOVA followed by Tukey’s HSD analyses were performed For exact *p* values see Dataset S1. (E) Representative confocal images correspond to (D). scale bars = 50 µm (F) kC9 Treatment to constitutively active *YDA* (*CA-YDA*) seedlings. Note that *CA-YDA* homozygotes (+/+) exhibit a pavement cell only phenotype, while heterozygous plants show an increase in small stomatal-lineage cells upon 50 µM kC9 treatment. (G) kC9 Treatment to dexamethasone-induced overexpression of NtMEK2^DD^, which is equivalent to MKK4/5 constitutive activation. (H) MAPK activation assays. Immunoblot analysis showing phosphorylated MAPKs (pMPK3/6) in wild-type seedlings treated with mock or 50 µM kC9 for 6 hrs. Here, the blots are overexposed to visualize the basal level of MAPK activation. Actin was used as a control. Ponceau S staining was used to ensure equal loading of protein samples. Normalized signal intensities relative to mock treatment are indicated below the blot.

We further dissected the MAPK pathway impacted by kC9. It is known that YDA MAPKKK and MKK4/5 MAPKK relay phosphorylation signals to MPK3/6 to enforce stomatal development (Herrmann and Torii, 2021). As such, a constitutively activation of YDA, CA-YDA, as well as that of MKK4/5 by a tobacco (*Nicotiana tabacum*) ortholog of Arabidopsis MKK4/5, NtMEK2^DD^, strongly activate MPK3/6 and confer astomatous epidermis (Bergmann et al., 2004; Wang et al., 2007). Treatment of kC9 exerts no effects on this ‘pavement cell only’ phenotype of CA-YDA and induced NtMEK2^DD^ overexpression (Fig. 2F, G), indicating that the constitutive activation of the YDA-MKK4/5-MPK3/6 pathway prevents kC9 from exerting its effects. Consistent with the notion that CA-YDA acts in a dose-dependent manner due to a weaker MAPK activation (Lampard et al., 2008; Wang et al., 2007), heterozygous CA-YDA plants responded to kC9 treatment with increased small, stomatal-lineage cells (Figs. 2F, S2D). On the other hand, weak co-suppression lines of *YDA* showed diminished efficacy of kC9 (Fig. S2E, F). A single loss-of-function *mpk3* and *mpk6* mutant, however, still responds to kC9 treatment, consistent with their functional redundancy (Fig. S2G, H).

These mechanistic dissections of kC9 action within the EPF-ERECTA-family signaling pathway narrow down the kC9’s *in vivo* targets to MAPKs. To directly address this possibility, we performed an established, immunoblot-based MAPK activation assays using anti-active MAPK antibody (Saijo et al., 2009). As shown in Fig. 2H, treatment of Arabidopsis wild-type seedlings under a stationary condition with 50 μM kC9 significantly reduced the activated population of MPK3/6 *in vivo*. Taken together, we conclude that kC9 triggers excessive stomatal differentiation via inhibiting the MAPKs, whereas a constitutive activation of the MAPK cascade incapacitates the action of kC9.

### kC9 directly binds to MPK6 and inhibits kinase activity

To decipher the molecular action of kC9, we first investigated whether kC9 inhibits MAPK activity via direct binding. For these purposes, we chose MPK6 for its available structural information (Putarjunan et al., 2019). As shown in Fig. 3A, *in vitro* kinase activity assays show that kC9 (**2**) inhibits the activity of recombinant MPK6 in a dose-dependent manner in a slightly weaker extent to a known MAPK activation inhibitor, U0126 (Favata et al., 1998). In contrast, the inactive analog kC9-3 (**3**) has no such effect. Next, we examined the *in vitro* direct binding assays of kC9 (**2**) and recombinant MPK6 using Biolayer Interferometry (BLI) and isothermal titration calorimetry (ITC) assays. kC9 (**2**) showed binding to MPK6 with a Kd of 4.9 ± 0.6 μM. In contrast, the inactive analog kC9-3 (**1**) did not exhibit any significant binding (Fig. 3B, Fig. S3, Tables S1, S2). Because the activation of the MAPK cascade incapacitates kC9 from increasing the number of stomata (Fig. 2, Fig. S2), we predicted that the constitutive activation of MPK6 may hinder kC9 binding. Indeed, a constitutively active CA-MPK6, MPK6 _D218G, E222A_, showed severely-reduced binding to kC9 with a Kd value of 58.0 ± 13 μM (Fig. 3B, Fig S3, Tables S1, S2). In addition, a kinase inactive DN-MPK6, MPK6 _K92M, K93R_, mutation slightly decreased the kC9 binding (Fig. S3, Tables S1, S2).

**Figure 3.**
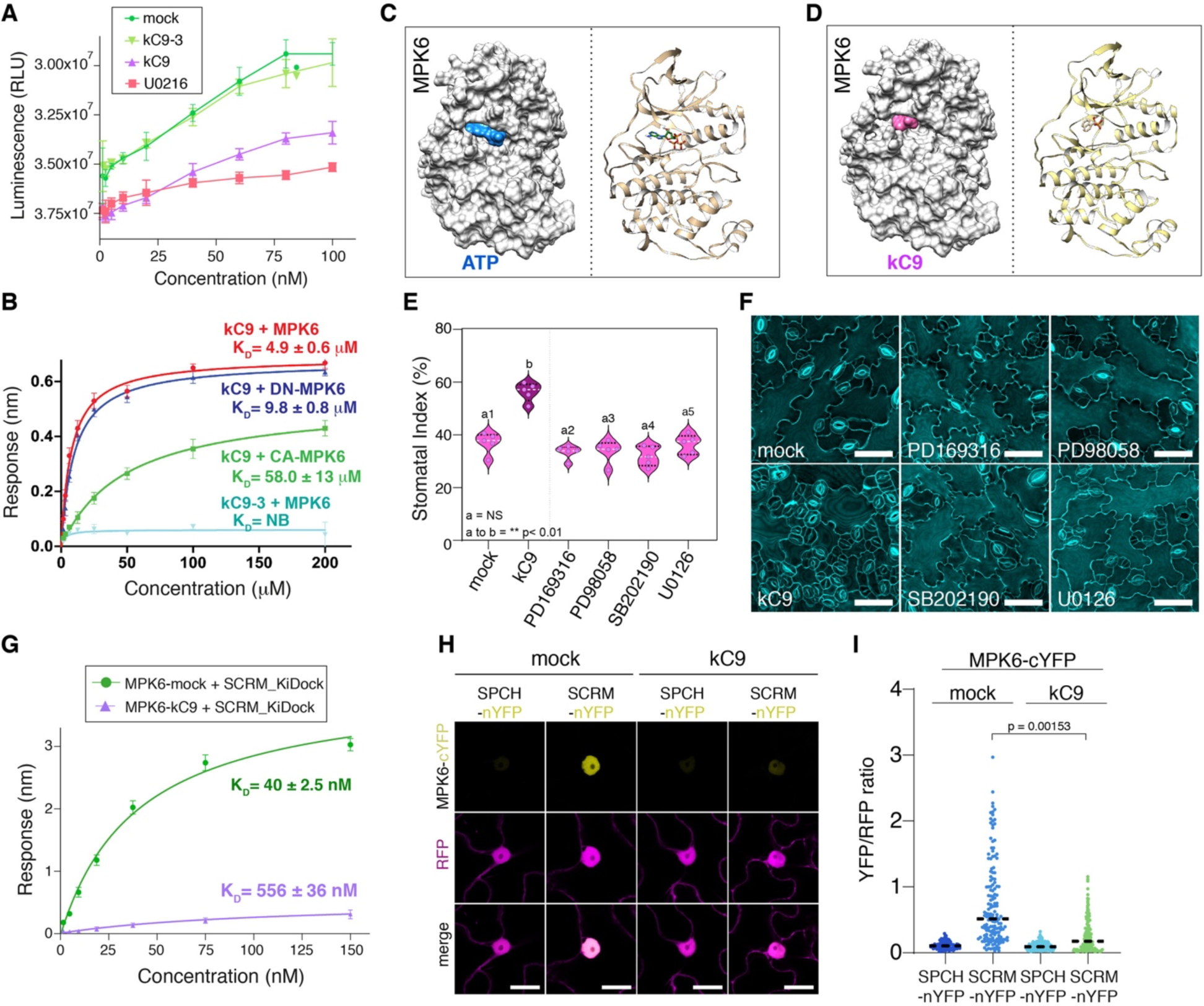
kC9 inhibits MPK6 activity and interfere with the MPK6-SCRM association. (A) Kinase activity assays for MPK6 in a concentration dependent manner in the presence of mock (dark green), 10 µM kC9-3 (light green), 10 µM kC9 (purple) and 10 µM of the animal MAPK inhibitor U02196 (light red). A decrease in luminescence (RLU) indicate MAPK activity. While mock and kC9-3 treatment does not affect MPK6 activity, both kC9 and U02196 show high luminescence after application, indicative of their inhibitory effects on MPK6 activity. Y-Axis has been inverted for visualization. (B) Quantitative analysis of kC9 binding kinetics to MPK6 as well as its inactive MPK6_K92M/K93R_ (DN-MPK6) and constitutively active MPK6_D218G/E222A_ (CA-MPK6) versions. The wild-type MPK6 exhibits higher affinity to kC9, while kC9-3 does not show any significant binding with MPK6. *In vitro* binding response curves are provided for kC9 with MPK6, MPK6_K92M/K93R_, and MPK6_D218G/E222A_, as well as for kC9-3 with MPK6. The data represent the mean ± SD and are representative of two independent experiments. (C, D) Structural docking models of known MPK6 structure (PDB: 6TDL) with ATP (A; blue) and kC9 (B; magenta) sharing a similar binding site. (E). Stomatal index of seven-day-old abaxial wildtype epidermis treated with different MAP Kinase inhibitors (mock, 50µM kC9, 10µM PD69316, 10 µM PD98058, 10 µM SB202190, 10 µM U01296) (E). Note that there is no significant difference of the stomatal index when treated with animal MAP Kinase inhibitors (a2-a5) compared to mock (a1) controls, while kC9 (b) increases the stomatal index significantly (a to b, **p = 0.0010053). 1-way ANOVA followed by Tukey’s HSD analyses were performed. For exact *p* values see Dataset S1. (F) Representative images of seven-day-old abaxial wildtype epidermis stained with PI and treated with different MAP Kinase inhibitors. Corresponds to (E). Scale bar = 50 µm. (G) Quantitative analysis of MPK6 with SCRM_KDock peptide interactions in the presence and absence of kC9. In vitro binding response curves for MPK6 and MPK6 with kC9 (1:2) with biotinylated SCRM_KDock peptide at seven different concentrations (150, 75, 37.5, 18.7, 9.3, 4.6, 2.3 nM) are subjected to analysis. Data are presented as mean ± SD, representative of three independent experiments. (H, I) Ratiometric bimolecular fluorescence complementation (rBiFC) analysis of interaction between MPK6 and SCRM as well as SPCH (negative control) in the absence (mock) and presence of kC9 assayed in *N. benthamiana* epidermis leaves. Images show maximum projections of n/cYFP fluorescence in nuclei, RFP fluorescence (internal control) and merge images (H). Bar = 20 µm. (I) Graph of YFP/RFP fluorescence ratios for protein pairs as indicated. For each protein pair single optical sections of nuclei, n/cYFP fluorescence and co-expressed cytosolic RFP serving as an internal control have been imaged. Mean n/cYFP and RFP fluorescence intensities were determined, and the ratio calculated for each nucleus. No or less efficient interaction results in a low YFP/RFP ratio (MPK6-cYFP+SPCH-nYFP, n = 126), whereas efficient n/cYFP complementation because of protein interaction results in elevated YFP/RFP ratios. Note that the efficiency of n/cYFP complementation for MPK6-cYFP + SCRM-nYFP (n = 158), is significantly reduced in the presence of kC9 compared to mock control. 1-way ANOVA followed by Tukey’s HSD analyses were performed. For exact *p* values see Dataset S1.

The reduced binding of kC9 to CA-MPK6 (and to some extent to DN-MPK6) implies that activation-loop dynamics play a prominent role in kC9 binding, suggestive of its binding to the ATP-binding pocket of MPK6. To address this, we took both experimental and docking simulation approaches. First, we observed that kC9 can effectively compete with ATP for binding to MPK6 (Fig. S3). The ATP binding affinity decreased approximately 2.5-fold in the presence of kC9 using ITC, indicating that either kC9 competes directly with ATP at the binding site or both share a similar binding pocket. Second, we conducted docking modeling studies to identify the specific binding pockets for kC9 and ATP. Briefly, we utilized HADDOCK-based calculations to guide the docking process, incorporating binding site information from Swiss Dock as ambiguous interaction restraints (de Vries et al., 2007; Dominguez et al., 2003). The resulting structural models indicate that kC9 fits to the ATP-binding pocket the binding interfaces involve a combination of hydrophobic and charged residues (Fig. 3C, D, Fig. S3). Due to single phosphate group in kC9 can form only two hydrogen bonds with MPK6 compared to five between ATP and MPK6 (Fig. S3). This weak binding mode may be reflected by the nature of kC9 not to impact stomatal development when the MAPK cascade is activated (Fig. 2, Fig. S2). In contrast, the inactive analog kC9-3 (**1**) as well as kC9-8 docking with MPK6 generated very few high energy structures, and these ligands are transiently fit at the narrow ATP binding pocket between N-and C lobes due to their bulky nature compared to kC9 (Fig. S3, Table S3). The docking results suggest that kC9 is a more favorable ligand compared to its inactive counterparts at the ATP binding pocket. Based on these findings, we conclude that kC9 directly inhibits MAPK (MPK6) via competitively binding to its ATP-binding pocket and that kC9’s binding is perturbed when MPK6 is activated.

### kC9 interferes with MPK6 - SCRM binding

Many small molecule inhibitors of MAPKs have been identified and utilized pharmacologically. Having identified kC9 as a direct inhibitor of MPK6, we sought to explore its selectivity and effectiveness for stomatal development. To address the question, we first compared the effects of known kinase inhibitors on stomatal development (Fig. 3E, F; see Methods). Unlike kC9, none of the known MAPK inhibitors we have tested (PD169316, PD98058, SB202190, and U0126 (Baltensperger et al., 1992; Gallagher et al., 1997; Liu et al., 2001) triggered an increase in SI or resulted in severe clustering albeit (Fig. 3F, G). Some inhibitors (SB202190 and U0126) caused aberrant cell division which resulted in guard cell pairs (Fig. 3F). However, these are not nearly as severe as stomatal cluster formation by kC9, suggesting that kC9 is highly potent and specific in stomatal development.

It has been shown that MPK6 is directly recruited to the SPCH-SCRM heterodimers via a bipartite MAPK binding motif within SCRM, named a KiDoK motif (Putarjunan et al., 2019). The mutation within this motif disrupts MPK6-SCRM interaction, thereby stabilizing the SPCH-SCRM heterodimers and resulting in a ‘stomata-only’ epidermis (Kanaoka et al., 2008; Putarjunan et al., 2019). The predicted MPK6-KiDoK binding interface (Putarjunan et al., 2019) partially overlap with the kC9-binding grove. We thus postulated if kC9 perturbs MPK6-KiDOK (hence MPK6-SCRM) association in addition to inhibiting the kinase activity. To test this hypothesis, we first performed BLI and ITC. Preincubating kC9 with MPK6 decreased the binding affinity of MPK6 and the SCRM KiDoK domain by 80-100% for both BLI and ITC (Kd values increased from 40.1 ± 1.2 nM to 856.5 ± 16.0 nM in BLI and from 43.4 ± 2.1 nM and 1403 ± 40 nM in ITC with or without pre-incubation with kC9, respectively) (Figs. 3G, Fig. S3L, M, Tables S1, S2). We further examined if kC9 perturbs the MPK-SCRM association *in planta* by ratiometric bimolecular fluorescent complementation (rBiFC) assays. In this setup, MPK6 and SCRM (or control SPCH), each fused with a complementary half YFP, along with a full-length RFP for normalization are located on the same plasmid, which were co-expressed in *Nicotiana benthamiana* leaves (see Methods). A strong YFP signal was reconstituted in MPK6-cYFP and SCRM-nYFP, confirming their strong direct association (Figs. 3H). Strikingly, kC9 treatment significantly diminished the YFP signals (Fig. 3H, I). SPCH-nYFP served as a negative control as it has been shown that MPK6 does not directly associate with SPCH (Putarjunan et al., 2019) (Fig. 3H, I). Collectively, these results affirm that kC9 binding not only attenuates the kinase activity of MPK6 but also disrupts the association with its direct substrate SCRM. This dual mode of kC9 may underpin the propensity of kC9 to induce excessive stomatal cluster formation.

### kC9 attenuates immunity response

We have demonstrated that kC9 exaggerate the number of stomata via directly inhibiting MPK6 activity as well as perturbing the recruitment of its downstream target SCRM, the latter of which likely contributes to the high propensity of kC9 to impact stomatal development (Figs. 1-3, S1-3). It is well established that MPK6 mediates diverse abiotic (environmental) and biotic stress signaling pathways, including drought-, osmotic-, salt-, and temperature stresses as well as immunity/defense responses (Danquah et al., 2014; Meng and Zhang, 2013; Sun and Zhang, 2022; Xu and Zhang, 2015). To gain a comprehensive view of signaling pathways influenced by kC9, we took an unbiased approach and characterized transcriptome dynamics in response to kC9 treatment. Specifically, wild-type seedlings were subjected to mock- or kC9 treatment for 6 and 24 hours followed by RNA-sequencing (Fig. 4A-C, Dataset S2, see Methods). We identified 500 and 641 differentially expressed genes (DEGs) between mock vs. kC9 treatment for 6 hrs and 24 hrs, respectively with false discovery rate [FDR] <0.01. Of these DEGs, 228 genes overlapped both 6 hrs and 24 hrs. Thus, 913 genes were detected as DEGs between mock vs. kC9 (Fig. S4).

**Figure 4.**
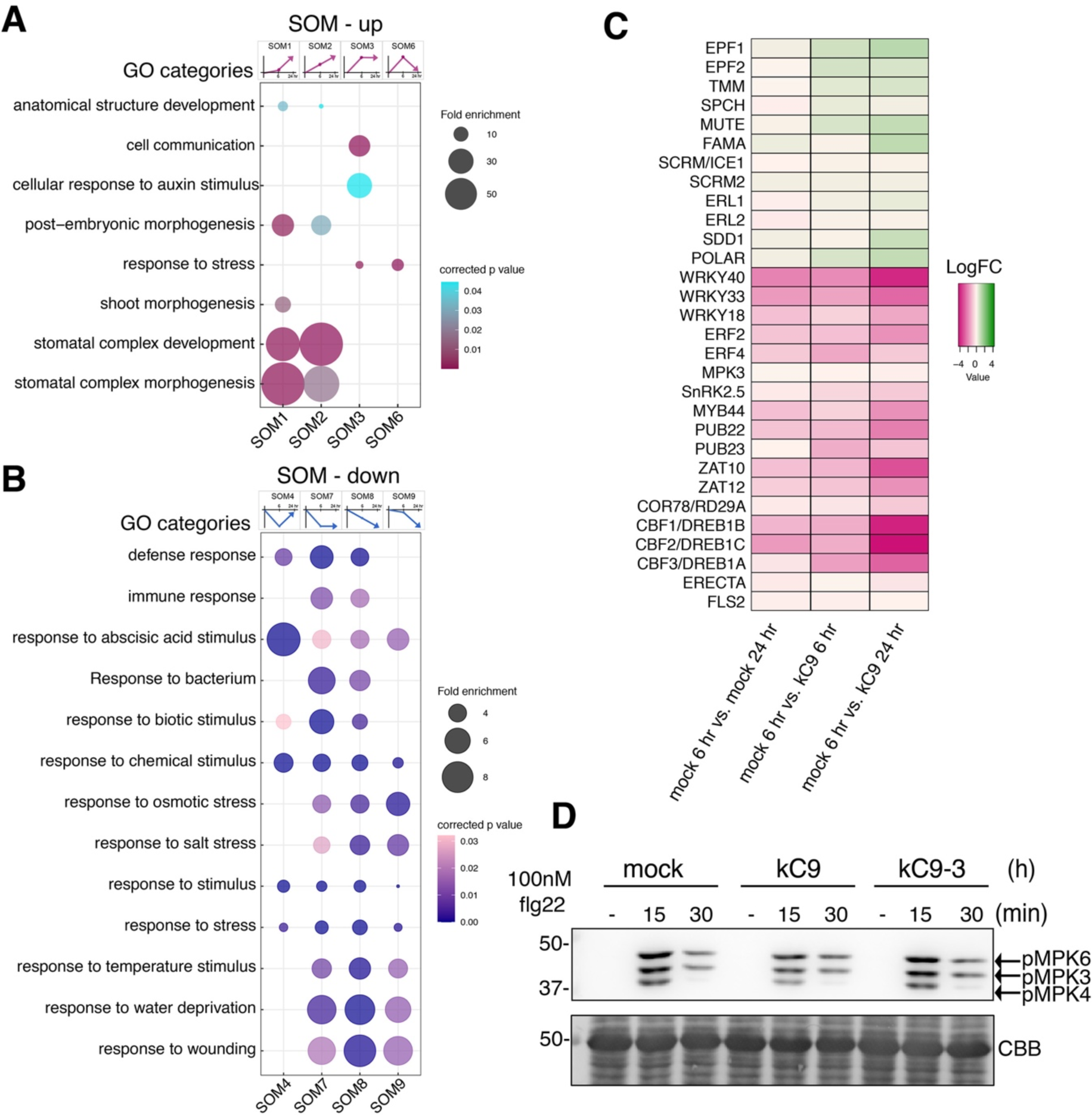
kC9 attenuates stress and immunity pathways. (A, B) Self-organizing map (SOM) clustering and GO enrichment of up-regulated (A) and down-regulated (B) differentially expressed genes (DEGs) categories. The gene expression trajectories of each SOM category are indicated as a diagram above the corresponding column of bubble plots. Selected, major GO terms are visualized by bubble plots. Fold enrichment and corrected p values of each cluster are visualized according to the bubble size and color scale, respectively. The number of genes in each SOM category: SOM1, 125; SOM2, 100; SOM3, 93; SOM4, 112; SOM6, 42; SOM7, 127; SOM8, 174; SOM9, 140. SOM cluster 5 is absent. See also Data S2. (C) Heat map showing the expression changes of the selected genes in upregulated (A) and downregulated (B) SOM clusters. Those from the upregulated SOMs include stomatal development genes, and those from the downregulated SOMs include biotic (e.g. immunity) and broad abiotic (cold, drought, ABA or ethylene mediated) pathways. The heatmaps present the mean log_2_(fold-change) values, relative to the corresponding levels in the mock 6-hour treatment. See also Data S2. (D) kC9 attenuates the MAPK activation in immune signaling. Top, immunoblot by anti-pERK antibody of seedlings treated with 100 nM flg22 for 0, 15, and 30 min in mock (left), kC9 (middle), or inactive kC9-3 (right). Bottom, total proteins stained with CBB. The locations of phosphorylated MAPKs are indicated by the arrows at the right. Molecular mass marker size is indicated at the left.

To categorize the dynamic response patterns of the DEGs, we next performed 3 x 3 self-organizing map (SOM) clustering based on the Log2(FC) values of the three-time points, including time 0, where theoretically no difference exists between mock and kC9. The SOM clusters were then classified into up- or down-regulated groups, after which gene expression (GO) categories overrepresented within each SOM were identified (see Methods). Consistent with the kC9’s effects, GO categories “stomatal complex development” and “stomatal complex morphogenesis” are highly enriched in the up-regulated SOM clusters, SOM1 (slowly up) and SOM2 (constantly up) (Fig. 4A). Other developmental processes “shoot morphogenesis”, “embryonic morphogenesis” and “anatomical structure development” are also detected in these SOMs. In SOM3 (up and stay steady) incudes “cell communication” (Fig. 4A). Consistently, stomatal regulatory genes are uniformly upregulated by kC9 treatment (Fig. 4C).

Strikingly, down-regulated SOM clusters are overwhelmingly dominated by abiotic and biotic stress response categories (Fig. 4B). Various biotic stress response categories, “defense response”, “response to biotic stimulus” are enriched in SOM4 (rapidly down and then back up), SOM7 (rapidly down and stay down), and SOM8 (constantly down). “Immune response” and “response to bacterium” are enriched in SOM7 and SOM8. Abiotic stress response categories are overrepresented in SOM7, SOM8, and SOM9 (slowly down then down), including “response to osmotic stress”, “response to salt stress”, “response to temperature stress”, “response to water deprivation”, and “response to wounding”. General stress response categories, such as “response to stimulus”, “response to stress” as well as “response to abscisic acid stimulus” were enriched across all four down-regulated SOMs (Fig. 4B). kC9 significantly repressed the expression of key TF genes, such as *CBF1/DREB1B*, *CBF2/DREB1C,* and *CBF3/DREB1A* for temperature/drought stress response as well as *WRKY33* and *WRKY40* for immunity response (Pandey and Somssich, 2009; Thomashow, 2001; Yamaguchi-Shinozaki and Shinozaki, 2006)(Fig. 4C).

To test if kC9 application attenuates the activity of MAPKs in the context of stress signaling, we turned our focus on a well-established immune signaling pathway (Asai et al., 2002). As previously demonstrated (Saijo et al., 2009), the application of flg22 peptide to Arabidopsis seedings elicits rapid phosphorylation and activation of MPKs (MPK3/6/4) within 15 min (Fig. 4D). Pre-incubation with kC9 slightly diminishes the flg22-induced phosphorylation, suggesting that kC9 can attenuate the activation of immune signaling by flg22 (Fig. 4D). Collectively, our results suggest that kC9’s potency to inhibit MPK6 extends beyond the context of stomatal development to diverse stress responses, including immune signaling.

### Activation of immune signaling nullifies kC9 from inducing stomatal development

Signal transduction pathways regulating immune response and stomatal development are initiated upon the recognition of distinct ligands (EPFs vs. flg22) by cognate primary receptors (ERECTA-family vs. FLS2)(Chinchilla et al., 2007; Lee et al., 2012). Yet, many of their signaling components are shared (Chen and Torii, 2023) (Fig. 5A). This raises an important, unresolved question of how their signal specificity can be maintained. We identified kC9 as a direct MPK6 inhibitor impacting stomatal development (Figs. 1-3). Moreover, kC9 potentially attenuates the flg22-mediated MAPK activation (Fig. 4). This prompted us to explore kC9 as a molecular tool to dissect the signal specificity.

**Figure 5.**
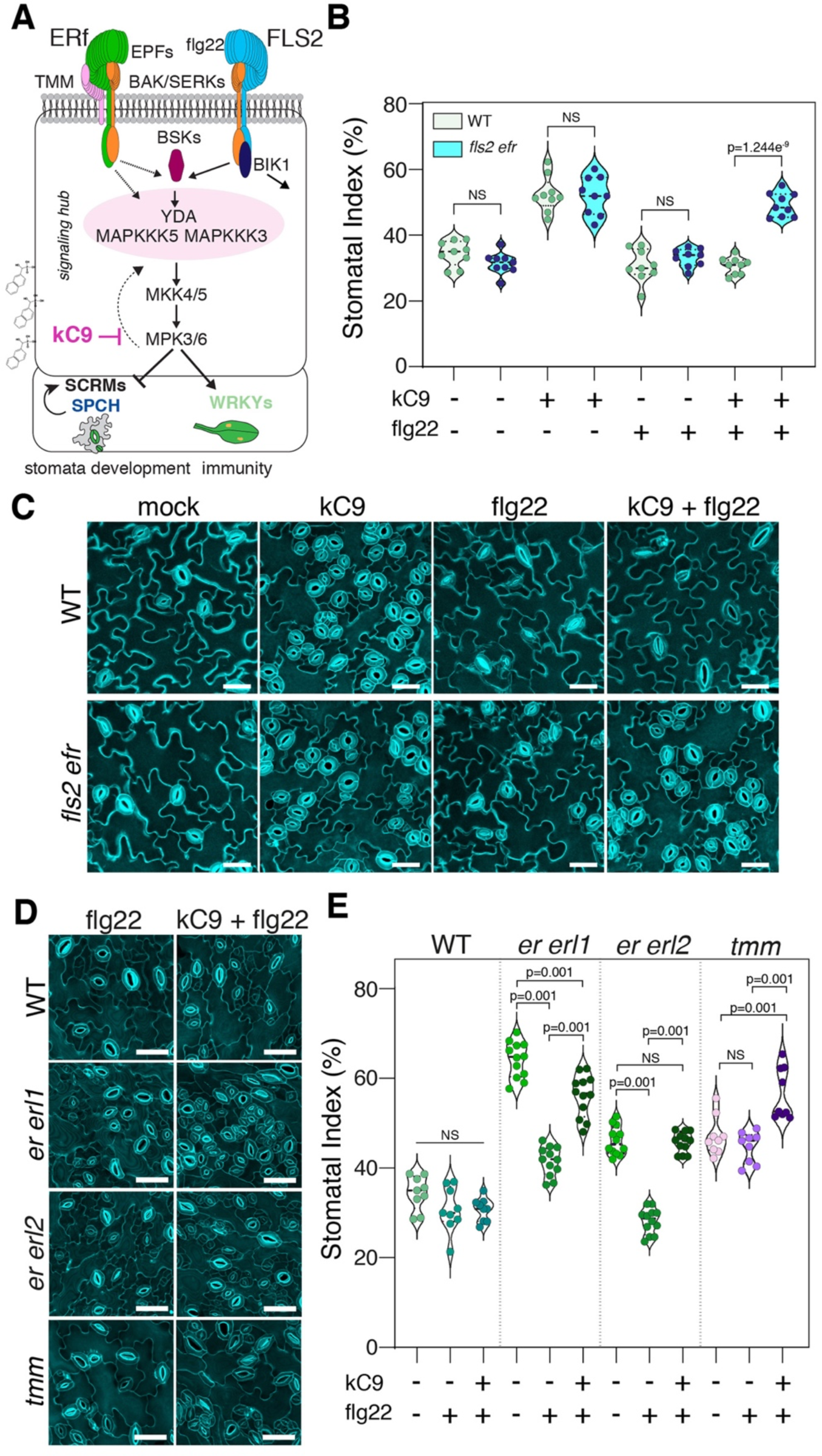
Activation of immune signaling nullifies excessive stomata formation by kC9 or *erecta*-family mutations. (A) Schematic representation of both the developmental and immune pathway (modified from Herrmann and Torii 2021). Note that both pathways do share signalling components. (B) Stomatal index (SI) of WT (n = 9) compared to *fls2 efr* (n = 9) double mutant seedlings treated with either mock (NS: 0.0901), kC9 (NS: 0.8927), flg22 (NS: 0.2065) or kC9 + flg22 (****p = 1.2435 E-09). Note that flg22 treatment can nullify kC9’s effect in WT but not *fls2 efr* double mutant plants. Two-tailed unpaired Student’s t-test was performed. (C) Representative confocal images of seven-day-old abaxial cotyledon epidermis from WT (top) and *fls2 efr* seedlings (bottom) correspond to (B). Bar = 40 µm (D, E) Activation of immunity pathway by flg22 reduces the stomatal index of stomatal development receptor mutants while co-treatment with kC9 can reverse the effect. (D) Representative confocal images of seven-day-old abaxial cotyledon epidermis from WT, *er erl1*, *er erl2* double mutant and *tmm* single mutant seedlings (from top to bottom) treated either with flg22 (left panel) or kC9 + flg22 (right panel). Bar = 50 µm. (E) Stomatal index of WT (n = 9) and stomatal component mutants (*er erl1*; n = 12, *er erl2*; n = 12, *tmm*; n = 9, from left to right) treated with either mock, flg22 or kC9 + flg22. The SI in WT does not show a significant change when either treated with flg22 or kC9 + flg22 compared to mock. In contrast, the SI is significantly reduced (**p < 0.001) in both *er erl1* and *er erl2* double mutant plants upon flg22 application. kC9 + flg22 co-application significantly increases (**p < 0.001) the SI for all three receptor mutants. 1-way ANOVA followed by Tukey’s HSD analyses were performed. For exact *p* values see Dataset S1.

kC9 application did not affect the expression levels of *ERECTA* and *FLS2*, the main receptors for stomatal development and innate-immunity response, respectively (Gomez-Gomez and Boller, 2000; Shpak et al., 2005) (Fig. 4C) To address if signal transduction pathways that enforce stomatal development are safeguarded from the activation of MPK6 by other (e.g. immunity) pathways, we conducted a separate and co-treatment of kC9 and flg22 peptide on wild-type seedlings. As previously demonstrated (Hohmann et al., 2020), treatment of flg22 alone does not affect stomatal development (Figs. 5B, C). Strikingly, however, the flg22 treatment completely blocked the excessive stomatal development triggered by kC9, resulting in the SI being insignificant from the mock-treated seedlings (Fig. 5B, C). The effect of flg22 to nullify kC9 is dependent on the presence of the corresponding immune receptor: *fls2 efr* mutant seedlings, which lack the major immune receptors FLS2 and EFR, are capable of fully responding to kC9 to increase stomatal development regardless of the presence or absence of flg22 (Fig. 5B, C). These results indicate that activation of the flg22-FLS2 immune signaling pathway can fully nullify kC9’s action as an MPK6 inhibitor triggering excessive stomatal development. These findings argue against the mechanism that separates MAPK cascades activated by different ligand-receptor pairs in order to safeguard signal specificity.

### flg22-mediated signal activation mitigates the loss of ERECTA-family receptors

We observed that flg22 application nullifies the effects of kC9 (Fig. 5B, C). Because kC9 shows reduced binding to CA-MPK6 (Fig. 3), it is plausible that kC9 cannot effectively bind to and inhibit the MPK6 (and likely MPK3) activated by flg22 treatment. If so, how can a stomatal cluster phenotype be ‘rescued’? It could be possible if the activated FLS2 signaling penetrates into the stomatal development pathway in the presence of kC9. This raises a question of whether this ‘merger’ of signaling pathways occur due to the artificial influence of chemical compounds or an inherent attribution of the endogenous pathways. To address this question, we examined the effects of flg22-induced immune signal activation on single and double loss-of-function *erecta*-family mutants that are partially compromised in MAPK activation (Lampard et al., 2008) (Figs. 5D, E, S5). As previously reported (Shpak et al., 2005), single loss-of-function mutant of *erl1* and *erl2* do not exhibit any stomatal phenotypes. However, we found that these single mutations attenuate the efficacy of the flg22 peptide in nullifying the elevated SI by kC9 (Fig. S5). This trend becomes more exaggerated in *er* single as well as *er erl1* and *er erl2* double mutants, where flg22 application no longer fully counteracts kC9 (Figs. 5D, E, S5).

Under normal (mock) conditions, these higher-order mutant seedlings exhibit higher SI than wild type owing to reduced receptor populations (Fig. S5). Surprisingly, even in the absence of kC9, the application of flg22 peptide alone is sufficient to inhibit the elevated SI in *er*, *er erl1*, and *er erl2* mutant seedlings (Figs. 5E, S5). These effects were less pronounced in the *tmm* mutant, perhaps owing to its axillary role for functional homeostasis of EPF-ERECTA-family signaling. Taken together, we conclude that under the reduced population of upstream ERECTA-family receptors, the activation of MAPK cascade by the flg22-mediated immunity pathway is sufficient to compensate for the reduced MAPK activation in the EPF-ERECTA-family stomatal development signaling pathways. The findings suggest that maintaining the specificity of signaling pathways relies on a proper homeostasis of available MAPKs that are yet to be activated.

### A vulnerability of stomatal-lineage cells to the immune signaling

Our study revealed that kC9’s effects in elevating stomatal numbers by inhibiting MPK6 activity can be counteracted by the activation of the MAPK cascade by the flg22-FLS2 immune signaling pathway (Fig. 5, S5). Moreover, even in the absence of kC9, flg22 can mitigate the excessive stomatal development due to reduced populations of the upstream ERECTA-family receptors (Fig. 5, S5). These findings challenge the prevailing notion of signal specificity (Ma and Nicolet, 2023). However, it is important to note that our experimental set-ups thus far entail a constant treatment of kC9 and/or flg22, therefore not fully accounting for the cellular response within the developmental timeframe of germinating seedling epidermis.

To delineate a developmental window where the altered MAPK activity by kC9 and/or flg22 impact stomatal differentiation, we designed a series of experiments as depicted in Fig. 6A. Here, either kC9 (*Experimental Design I*) or flg22 (*Experimental Design II*) were applied consecutively on each day after germination up to Day 7. In the co-treatment experiment (*Experimental Design III*), seedlings were germinated in the constant presence of kC9, with flg22 peptide sequentially applied on each day after germination (Fig. 6A). Cotyledon abaxial epidermis was observed at Day 7 (Figs. 6B, D, F, S6A-C). In *Experimental Design I*, the number of stomata declined as shorter the duration of kC9 treatment (Figs. 6B, C, S6A). The number of epidermal pavement cells stayed constant whereas that of small stomatal-lineage cells slowly decreased as stomata differentiate (Figs. 6E, G, S6A). Thus, the number of stomata inversely correlated with the duration and early application of kC9 treatment.

**Figure 6.**
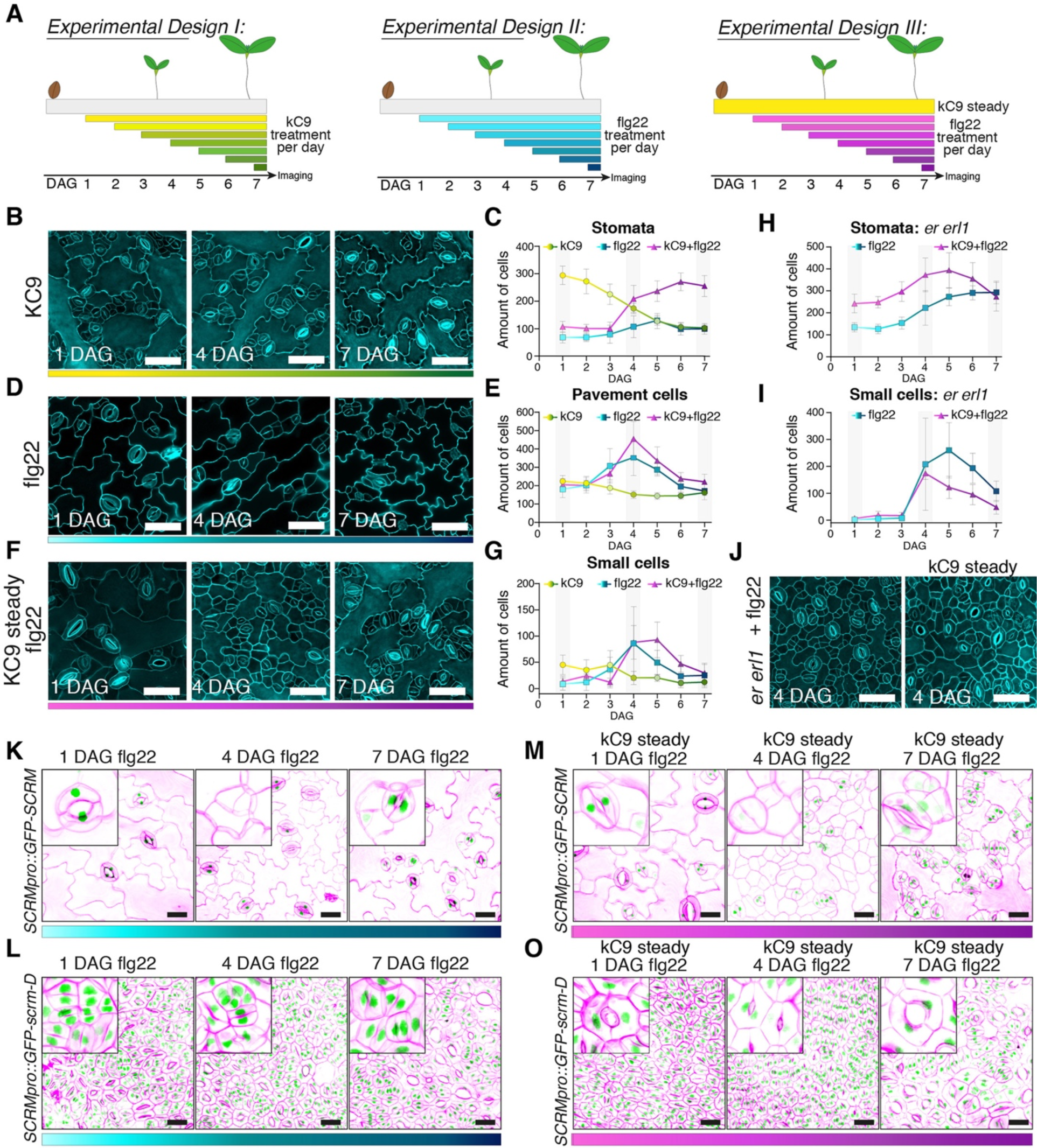
Sequential flg22 application reveals the developmental stage labile to the immune signal activation. (A) Schematics of time-course experiments. (left) kC9 treatment (*Experimental Design I*, yellow to green), (middle) flg22 treatment (*Experimental Design II,* light blue to dark blue), (right) flg22 treatment in the steady level of kC9 (*Experimental Design III*, pink to purple). (B-G) Chemical and/or peptide treatment over time depicted in (A) of wild type (WT) cotyledon abaxial seedlings epidermis. (B, D, F) Representative confocal images of seven-day-old WT abaxial cotyledon epidermis from seedlings treated on 1 DAG, 4 DAG, or 7 DAG with 50 µM kC9 (B, yellow to green), 0.1 µM flg22 (D, light blue to dark blue) or 0.1µM flg22 where kC9 steadily was present for 7 days (F, pink to purple). Note an accumulation of small epidermal cells in (F) on 4 DAG. Bars = 50 µm (C, E, G) Quantitative analysis of different epidermal cell types of WT cotyledon abaxial seedlings treated over time. Correspond to (A, B, D, F). For exact values see Dataset S1. For all graphs, color codes are: 50 µM kC9 (yellow to green); 0.1 µM flg22 (light blue to dark blue); 0.1µM flg22 where kC9 steadily was present for 7 days (pink to purple). Number of stomata (C), epidermal pavement cells (E) and small cells representing meristemoids and SLGCs (G) per unit area (0.339178112 mm^2^) are plotted. Error bars, S.D. Note that both flg22 application in the present or absent of kC9 results in a peak when applied on 4 DAG and 5 DAG (kC9 + flg22). (H, J) Chemical and/or peptide treatment over time depicted in (A) of seven-day-old *er erl1* double mutant cotyledon abaxial seedlings epidermis. (H, J) Quantitative analysis of different epidermal cell types in *er erl1* cotyledon abaxial seedlings treated over time on *Experimental Design II* (light blue to dark blue) and *III* (pink to purple). Number of stomata (H) and small cells representing meristemoids and SLGCs (I) per unit area (0.339178112 mm^2^) are plotted. Error bars, S. D. Note that in (H) flg22’s effect to suppress stomatal development is revertedin the present of kC9. Application of flg22 in the present or absent of kC9 results in a peak when applied on 4 DAG and 5 DAG. (J) Representative confocal images of seven-days-old *er erl1* cotyledon abaxial seedlings epidermis treated on 4 DAG with 0.1 µM flg22 in the present or absence of kC9. Note an accumulation of small epidermal cells in both. Bars = 50 µm (K-O) Representative confocal images of seven-days-old cotyledon expressing *SCRMpro::GFP-SCRM* (K, M) or *SCRMpro::GFP-scrm-D* (L, O) treated on 1, 4 or 7 DAG with 0.1 µM flg22 in the present (M, O) or absence (K, L) of kC9. Note that GFP-SCRM is not present in small cells when flg22 was applied on 4 DAG present (M) or absence (K) of kC9. On the contrary, application of flg22 on any given day does not affect accumulation of GFP-scrm-D (L, O). Bars = 25 µm.

In contrast, time window-specific effects of flg22 on stomatal development was observed when we performed *Experimental Design II*, a sequential application of the flg22 peptide (Figs. 6C, D, E, G, S6B). Whereas the number of stomata remained relatively constant throughout the time course, small stomatal-lineage-like cells become overwhelmingly increased, peaking at 4 DAG (4 days of flg22 treatment) (Figs. 6C, D, G, S6B). The vast increase of the small stomatal-lineage-like cells (and consequently pavement cells) is more notable in *Experimental Design III,* where flg22 was sequentially applied under the constant presence of kC9, again peaking the numbers at 4-5 DAG (4 to 3 days of flg22 treatment) (Figs. 6E-G, S6C). In the *er erl1* double mutant background whereby ERECTA-family receptor signaling is suboptimal, the effects of flg22 to vastly increase the small cells at 4-5 DAG became even more exaggerated (Fig. 6H, I).

To further explore the identity of these small cells induced by the flg22 application, we examined the expression of SCRM protein, a SPCH/MUTE/FAMA partner that is directly targeted by MPK6 (Kanaoka et al., 2008; Putarjunan et al., 2019). For this purpose, seedlings expressing a fusion protein of GFP and SCRM (*SCRMpro::GFP-SCRM*) and its stabilized version (*SCRMpro::GFP-scrm-D*) each driven by the endogenous promoter were subjected to *Experimental Designs I* and *III* (Figs. 6A, K-O). SCRM is known to accumulate the nucleus of all stomatal-lineage cells, including SLGCs (Kanaoka et al., 2008). We found that most of all massively accumulated small cells at 4 DAG do not exhibit GFP signals (Fig. 6K, L, middle panels). Likewise, transcriptional reporters of stomatal precursor cells, *SCRMpro::nucGFP*, *MUTEpro::nucYFP* and *TMMpro::GUS-GFP* are barely detectable when flg22 was applied at 4 DAG (Fig. S6D-I), indicating that these small cells lost the proper stomatal precursor identity. In contrast, flg22 has no effects on GFP-scrm-D accumulation nor ‘stomata-only’ phenotype conferred by GFP-scrm-D (Fig. 6L, O). Collectively, these results suggest that flg22 treatment at 4 DAG, when stomatal-lineage cells are actively proliferating (Pillitteri et al., 2011), triggers the arrest and subsequent withdrawal of stomatal-lineage fate through the MPK6-mediated degradation of SCRM. Moreover, the remarkably narrow developmental window of the flg22 action implies that stomatal-lineage cells are most sensitive to the activation of MAPK cascade by otherwise unrelated immune signaling pathways when the cells are in the decision-making process. This vulnerability likely stems from the intricate cellular response to a proper activation status of MAPKs.

## DISCUSSION

Through a small compound library screening, we identified kC9 as a potent inducer of stomatal development via inhibiting the canonical EPF-ERECTA-family signaling pathway. Further comprehensive chemical genetics analyses together with *in vitro* biochemical, biophysical and structural studies revealed that kC9 is a new inhibitor of plant MAPKs. Surprisingly, we found that the activation of immune RK signaling pathway fully nullifies the excessive stomatal differentiation by kC9. This cross-activation is not an artifact of chemical application, but rather, the signal integration results if the activable upstream receptors for stomatal development are limited. Finally, we revealed the cell-state specificity upon which the immune signaling pathway intervene stomatal differentiation through the activation of a shared MAPK cascade.

Our docking modeling of kC9 with the structurally solved MPK6 and subsequent binding and kinase activity assays, backed up by the structure-activity relationship analysis of the kC9 analogs revealed that kC9 weakly binds to the ATP-binding pocket of MPK6 and inhibits its activity (Figs. 3, S1, S3). The ATP binding pocket within the catalytic cleft is shared by protein kinases (Liao, 2007), as such kC9 may not likely be a highly-selective kinase inhibitor. Aligning with this view, kC9 (HNMPA) is a known insulin receptor tyrosine kinase (RTK) inhibitor (Baltensperger et al., 1992). Nevertheless, as demonstrated by the extensive phenotypic analyses and the *in vivo* MPK3/6 activity assays (Figs. 1, 2, S2), it is clear that kC9 inhibits MAPK activation downstream of the EPF-ERECTA-family. One explanation for the strong propensity of kC9 to impact stomatal development is that the binding of kC9 to MPK6 perturbs the recruitment of its substrate, SCRM (Fig. 3G-I). SCRM possesses a unique bipartite anchoring module (KiDoK, canonical Kinase docking site and SCRM-specific KRAAM motif) to associate with MPK6, and it is required for the post-translational regulation of SPCH and, hence, proper stomatal patterning (Putarjunan et al., 2019). MPK6 phosphorylates diverse regulators of development and environmental responses, including MYC2 in jasmonic acid response, WRKYs in immunity and pollen development (Guan et al., 2014; Sethi et al., 2014; Sun and Zhang, 2022). Although the structural basis of MPK6 binding mode with these substrates is not known, further studies may unravel exactly how kC9 impact additional MPK6-mediated pathways, as suggested by our transcriptome analysis (Fig. 4).

The diminished binding affinity of kC9 to the pre-activated MPK6 (Fig. 3B) enabled us to investigate whether the pre-activation of the upstream signaling receptors can forestall kC9 from inhibiting the MPK6-mediated inhibition of stomatal development. Strikingly, the kC9’s action can be nullified not only by the *bona fide* ligands for ERECTA-family RKs, EPF1 and EPF2, but also by the immunity ligand flg22 for FLS2 (Figs. 2, 5). In the cell signaling field, it is widely established that extracellular signals perceived by the cell-surface receptors are relayed to a specific MAPK cascade via selective activation of scaffold proteins (Garrington and Johnson, 1999). The well-known example is yeast mating and osmolarity response, both of which are mediated by the shared MAPKKK, Ste11. There, the activated upstream signaling recruits distinct scaffold proteins, Ste5 and Pbs2 for mating and osmosis, respectively, to ensure signal specificity (Choi et al., 1994; Park et al., 2003; Posas and Saito, 1997). In plants, RACK1 (RECEPTOR FOR ACTIVATED KINASE C) functions as a scaffold protein that bridges heterotrimeric G-proteins to an MAPK cascade (Cheng et al., 2015). Yet, scaffold proteins that separate LRR-RK immunity vs. development signaling remains elusive (Ma and Nicolet, 2023).

Based on our discovery, we propose that signal specificity of the EPF-ERECTA-family pathway can be ensured by its own fully activable state. During epidermal development, ERECTA-family RKs are constantly detecting multiple EPF/EPFL peptides (Herrmann and Torii, 2021; Torii, 2012), which likely elicits optimal, continual activation of the downstream MAPK cascade. This signaling dynamics itself prevents the inappropriate signal bleed-through from other inputs. Our findings accord with a previous report that the constitutive activation of the FLS2 immune signaling failed to impact stomatal development, when ERECTA-family signaling is fully active (Hohmann et al., 2020). Moreover, our proposed mechanism reconciles the previously reported inconsistencies regarding the relationship between YODA and MAPKKK3/MAPKKK5 in development and immunity, whether they act together or antagonistically (Liu et al., 2022; Sun and Zhang, 2022; Wang et al., 2022). Liu et al. (2022) observed that flg22 peptide triggers excessive phosphorylation of MPK3/6 in *erecta-family* single and double mutant backgrounds and concluded that the immune signaling can be hyper-activated when the population of ERECTA-family receptors are limited. However, this could be the sign of immunity-to-development signal cross activation as observed in our study (see Figs. 5D, E, S4A, B). On the other hand, whether the EPF-ERECTA pathway can also cross-activate the immune signal remains unclear. It has been reported that ERECTA regulates plant disease resistance via YODA MAPK cascade (Godiard et al., 2003; Jorda et al., 2016; Sopena-Torres et al., 2018). Since ERECTA’s role in immunity does not seem to require EPF peptides (Jorda et al., 2016), it may involve different mechanisms, such as an indirect resistance through modulation of downstream gene expression that influence cell wall properties (Jorda et al., 2016; Sopena-Torres et al., 2018).

It is important to note that even under suboptimal ERECTA-family signaling pathway, the activated flg22-FLS2 signaling pathway can only reduce the numbers of stomata to the wild-type level (see Figs. 5D, E, S4A, B). In contrast, the induced EPF1 or EPF2 overexpression, both of which severely diminish the number of stomata even in the presence of kC9 (Fig. 2D, E). These observations imply that influence of the immune signal on the MAPK cascade in stomatal development is safeguarded by a functional MAPK homeostasis, which is regulated by the strengths of the signal input *(i.e.* ERECTA-family RKs). Consequently, the cross-activation of the MAPK cascade by immunity signal inputs cannot exceed the basal (normal) level. While the exact molecular nature of the threshold that limits cross-signal activation is unknown, it is tempting to hypothesize that the MAPK cascade components form a shared dynamic signaling hub (Fig. 5A). This hub can sense and associate with a partially compromised upstream receptor to allow cross-activation, reflecting how signaling input strength and MAPK homeostasis contribute to signal specificity

Our time-course experiments (Fig. 6) revealed the narrow developmental window of the flg22-induced inhibition of stomatal development, highlighting the vulnerability of proliferating stomatal-lineage cells to the immune signaling. These early stomatal-lineage cells undergo asymmetric cell divisions, where the differential MAPK activities will be translated into the cell-fate decision to become a meristemoid or a stomatal-lineage ground cell (Zhang et al., 2023). Consistently, MAP KINASE PHOSPHATASE1 (MPK1), which attenuates MAPK activity, influences stomatal differentiation during early proliferating precursor state (Tamnanloo et al., 2018). In addition to the EPF-ERECTA-family-mediated MAPK signaling, dynamically-assembled polarity proteins including BREAKING OF ASYMMETRY IN STOMATAL LINEAGES (BASL) differentiate the MAPK activity between the two daughter cells via positive feedbacks (Zhang et al., 2016; Zhang et al., 2015). As such, it is conceivable that additional spike of immune signal activation disturbs the intricacy of differential MAPK activity required for maintaining the meristemoid identity. The cross-activation mechanism between developmental and immune signals in stomatal-lineage cells may also contribute to optical number of stomata that are critical for the inhibition of pathogen entry and subsequent prevention of pathogen growth (Chen and Torii, 2023). Understanding how different cell types and cell states interpret and respond to development and developmental signals is fundamentally important future direction to gain insight into the dynamics and heterogeneity in cellular signal specificity and integration.

## ACKNOWLEDGEMENTS

We thank Dr. Yueling Zhang for *ami-YDA* lines, Dr. Cyril Zipfel for *fls2 efr* seeds, Dr. Liangliang Chen for the thoughtful discussion, and Rachel Cole and Maggie Magaw for help in plant care. This work was supported by MEXT Japan Society for Promotion of Science (JSPS) KAKENHI (JP16H01237, JP17H06476, JP19H00990) to K.U.T., ITbM Research Award to H.E., and NIH (R35GM144275) and NSF (IOS-2049642) to L.S. A.H. was in part supported by the Walter Benjamin Program, Deutsche Forschungsgemeinschaft (DFG; 447617898). K.U.T. acknowledges the support from the Howard Hughes Medical Institute and Johnson & Johnson Centennial Chair of Plant Cell Biology at UT Austin.

## AUTHOR CONTRIBUTIONS

Conceived the project, K.U.T.; Performed chemical-genetics screen and assays, A.H., H.E., H.K., A.N.; Biochemical/biophysical assays, K.M.S., P.B., J.L.; Structural modeling, K.M.S.; Chemical synthesis, identification, derivatization, and analyses, S.K., A.Z., and S.H.; transcriptome profiling and bioinformatics, N.U., S.K.; Data analysis and visualization, A.H., S.K., K.M.S., L.S., K.U.T.; Writing, K.U.T.; Manuscript editing, K.U.T., K.M.S., A.H., L.S. and manuscript approved by the all authors; Funding acquisition, K.U.T., H.E., L.S.

## LEAD CONTACT

Further information and requests for resources and reagents should be directed and will be fulfilled by the Lead Contact, Keiko U. Torii.

## MATERIALS AVAILABILITY

Plasmids and transgenic plants generated in this study will be available from the lead contact upon request. Chemical compounds synthesized in this work, especially those not commercially available, may be provided in limited quantities.

## DATA AND SOFTWARE AVAILABILITY

- This paper does not report original code.
- The RNA-seq data generated in this study are deposited to the DDBJ (DNA Data Bank of Japan) with an accession number PRJDB18465
- For software for bioinformatics, image-analysis, and biophysical analysis, see Key Resources Table
- Any additional information required to reanalyze the data reported in this paper is available either from the Texas Data Repository Datavarse (http://datavarse.tdl.org/) or from the lead contact upon request

## EXPERIMENTAL MODEL AND SUBJECT DETAILS

The Arabidopsis Columbia (Col) accession was used as wild type. The following mutants and reporter transgenic lines were reported previously: *er-105, erl1-2, erl2-1, er-105 erl1-2*, *er-105 erl2-1, erl1-2 erl2-1, er-105 erl1-2 erl2-1* (Shpak et al., 2004); *spch-3*, *fama*, and *E994* (Pillitteri et al., 2007); *mute-2* (Pillitteri et al., 2008); *MUTEpro::nucYFP* (Qi et al., 2017); *SCRMpro::GFP-SCRM* and *SCRMpro::GFP-scrm-D* (Kanaoka et al., 2008); *TMMpro::GUS-GFP* (Nadeau and Sack, 2002); *tmm-KO* (Hara et al., 2007), *epf1-1, epf2-1, Est::EPF1, Est::EPF2* (Lee et al., 2012); *mpk3, mpk6, NtMEK2_DD_* (Wang et al., 2007); *fls2 efr* (Zipfel et al., 2006); *bak1-5,serk1-1, serk2-1* (Meng et al., 2015); *CA-YDA* (Lukowitz et al., 2004), *amiYDA#1, amiYDA#2* (Sun et al., 2018), and also indicated in Key Resource Table. Seedlings and plants were grown in a long-day condition at 21 °C.

## METHOD DETAILS

### Chemical screening

For the initial screen, we used Arabidopsis E994 line that expresses stomatal guard cell-specific erGFP. Three E994 seeds were germinated in each well of 96-well microtiter plate (TL5003; True Line) supplemented with 95 µL of 1/2 MS media. At 1-day-post germination, 5 µL of individual chemical compounds (DMSO/media mixture with the ratio of 1:9 prepared from 10 mM master stock in 100% DMSO) from the ITbM chemical compound library (Ziadi et al., 2017) were added to each well at the final concentration of 50 μM. The seedlings were grown for 9 days at 140 rpm, 22 °C in continuous light (and subsequently GFP signals were examined under Zeiss SteREO Discovery V20 fluorescent microscope. The compounds that increased GFP+ stomata were subjected to a secondary screen to validate the reproducibility of the compounds to increase numbers of stomata. Those chemicals that passed the secondary screen were moved to a tertiary screen using seedlings expressing *TMMpro::GUS-GFP* (Nadeau and Sack, 2002), which marks stomatal-lineage cells. Confocal images of *TMMpro::GUS-GFP* abaxial cotyledon epidermis are subsequently analyzed using our pipeline: The original images were converted to black/white, and area mean intensity of black pixels, which correspond to GFP signals from three biological replicates was calculated and plotted. Those compounds that increased the area mean intensity signals of stomatal-lineage cells > 1.5 fold (p < 0.05) are classified as Stomatal Number Increasing Compounds.

### Chemical synthesis and NMR analysis

We synthesized kC9 (Hydroxy-2-naphthalenylmethylphosphonic acid: HNMPA)(**2**) and its derivatives from commercially available 2-naphthaldehyde, 1-pyrenecarboxaldehyde, and 2-bromomethyl naphthalene. Phosphonic acid diesters (**1**, **5**, **7**, and **9**) were prepared by nucleophilic addition and substitution reactions with dialkyl phosphites. Acetylation of a-hydroxyl group of **1** led to **3**. Lewis acid treatment of phosphonic acid diesters (**1**, **3**, **5**, **7**, and **9**) successfully provided phosphonic acids (**2**, **4**, **6** and **8**) and phosphonic acid monoester (**10**), respectively. The ^1^H, ^13^C(Gianakopoulos et al.), ^31^P{^1^H} NMR were recorded on a JEOL JNM-ECA500 (500 MHz for ^1^H, 125 MHz for ^13^C, 202 MHz for ^31^P) spectrometer. Chemical shifts were reported in ppm (δ), and coupling constants were reported in Hz. High resolution mass analyses (HRMS) were submitted to the Mass Spectrometry Laboratory (Molecular Structure Characterization Unit) at RIKEN. Thin-layer chromatography was performed on Merck 60 F254 precoated silica gel plates. For detailed chemical synthesis procedures as well as NMR spectra data, see Document S1.

### Chemical treatment

Compounds. CL1 (SC-58125, Cat. No. 70655) was purchased from Cayman Chemical Company. Bubblin ([4-(4-bromophenyl)-2-pyridin-3-yl-1,3-thiazole]; MS-6628) was purchased from Key-Organics. kC9 and its analogs were synthesized (see above section). kC9 and its phosphonic acid ester form (*P*-[[(Acetyloxy)methoxy]-2-naphthalenylmethyl]-bis[(acetyloxy)methyl] ester phosphonic acid: **1**) are also commercially available (abcam AB141566 and abcam141567, respectively). These compounds were diluted in DMSO and subjected for chemical treatment. For experiments in Figs. 1B, F and 5E, seedlings were incubated for 8 days and observed at day 9. For all other experiments, seedlings were incubated for 7 days and observed at day 7. For a dose-dependent analysis, kC9 was diluted to respective concentrations from 100 mM stock solution and subjected to a forementioned treatments. For phenotypic analyses, kC9 were applied to Arabidopsis mutants and reporter lines using 24-well microtiter plates (Life Science, Durham, USA, REF: 351147). For estradiol induction of EPF1 (*Est::EPF1*) and EPF2 (*Est::iEPF2*) overexpression, germinated seedlings were grown in the presence of 5 µM estradiol with or without 50 µM Kc9. flg22 peptide was either synthesized at the ITbM Peptide Synthesis Center or obtain from BioSynthesis (Lot No.: K1254-1). Seedlings were treated with 50-100 nM flg22 peptide in 1/2 x MS liquid media in the presence or absence of 50 µM kC9.

### Confocal microscopy

For confocal microscopy, cell peripheries of seedlings were visualized with propidium iodide (Sigma, P4170). Images were acquired using LSM800 (Zeiss) using a 20x lens. The GFP, YFP reporter and PI signals were detected at with excitation at 561 nm and 582-617 nm emission range, respectively. Raw data were collected with 1024 x 1024 pixel images. Alternatively, images were acquired using Stellaris-8 FALCON (Leica - Mannheim, Germany) using either a 40x or 63× oil lens for high-resolution imaging and 20x water lens for data acquisition. Signals were detected for the following conditions: GFP, excitation at 488 nm and emission from 490 to 546 nm; YFP, excitation at 514 nm and emission from 520 to 560 nm; and Propidium iodide, excitation at 561 nm and emission from 570 to 620 nm. Signals were visualized sequentially using separate HyD detectors (HyDX/HyDS). For qualitative image presentation, Fiji (ImageJ) as well as Adobe Photoshop CS6 was used to trim and uniformly adjust the contrast/brightness.

### Plasmid Construction

The following constructs were generated for the study: pAHs022, pAHs39, pAHs041, pAHs045, pAHs053, pAHs55. For detailed information of each plasmid, see Supplementary Table S4. See Table S5 for a list of primer oligo DNA sequence. For pDONR221_P3P2, coding sequence of SCRM as well as SPCH (Putarjunan et al., 2019) were amplified (Phusion, NEB, LOT:10187023) and cloned via a Gateway BP reaction (Thermo Fisher Scientific, LOT: 2536741) into *pDONOR221_P3P2*. Subsequently, Gateway LR reaction (Thermo Fisher Scientific, LOT: 2600344) was performed using *pBiFCt:2in1-CC* (256) as destination vector. For pDONR221_P1P4, MPK6 coding sequence (Putarjunan et al., 2019) was cloned via a Gateway BP reaction (Thermo Fisher Scientific, LOT: 2536741) into *pDONOR221_P1P4*. Subsequently, Gateway LR reaction (Thermo Fisher Scientific, LOT: 2600344) was performed using *pBiFCt:2in1-CC* (256) as destination vector. To create a dominant negative (DN, K92M_K93R) and constitutive active version (CA, D218G_E22A) of MPK6 for protein purification, a site directed mutagenesis was performed using the Prime STAR (Takara Bio, LOT: N5101FA) protocol and *pGEX4T-1_MPK6ΔNt (29-305)* (Putarjunan et al., 2019) as backbone.

### Ratiometric Biomolecular Fluorescence Complementation (rBiFC)

The Gateway compatible 2in1 system (Grefen and Blatt, 2012; Mehlhorn et al., 2018) was used to test interaction between SCRM and MPK6 in the presence or absence of Kc9. SPCH together with MPK6 was used as negative control. Amplicons with recombination sites were cloned either in pDONOR221-P3P2 or pDONOR221-P1P4 by BP clonase respectively. Subsequent LR clonase reaction was performed with the pBiFCt-2in1-CC destination vector. While both SPCH and SCRM were fused to nYFP, MPK6 was fused to cYFP on the same plasmid, each expressed under the control of a 35S promoter. An internal expression control (35S::RFP) is also included on the plasmid. Interaction strength was determined by the YFP/RFP ratio. For this, 4-week-old leaves of *N. benthamiana* were transfected with *A. tumefaciens*, carrying respective destination vectors at an OD of 0.2 in the presence or absence of 100µM Kc9 and incubated for two days in the plant room under short day conditions. Confocal microscopy single plane images of nuclei were taken by using the Stellaris-8 FALCON (Leica - Mannheim, Germany) and a 63× oil lens. For comparison of interaction strength rBiFC, identical confocal settings were used for respective experiments (n = 3, ∼ 50 nuclei each). The average fluorescence signal intensity for YFP and RFP was measured by using ImageJ v1.8.0_66. Calculation of mean YFP/RFP ratios were performed in Excel and graph was assembled in GraphPad Prism v10.

### RNA sequencing and analysis

Three-day-old Arabidopsis wild-type seedlings were treated with mock (5% DMSO) or 50 µM kC9 (in 5% DMSO) for 6 and 24 hours, and total RNAs were extracted and purified using RNeasy Plant Mini Kit (QIAGEN). Each RNA sample (three independent RNA samples for each treatment) was prepared from a pool of 30 whole seedlings. 3 µg of purified RNA was used for RNA-seq as described previously (Uchida et al., 2018). Briefly, after RNA integrity was confirmed by running the RNA samples on Agilent RNA 6000 Nano Chip (Agilent Technologies), RNA-seq libraries were prepared using Illumina TruSeq Stranded mRNA LT Sample kit. The resulting barcoded libraries were pooled and sequenced on Illumina NextSeq500 sequencing platform, and 75-bp single-end reads were obtained. Sequences of RNA-seq experiments were mapped on TAIR 10 genome using TopHat2 (ver. 2.1.0, https://github.com/infphilo/tophat) with default options, and the Illumina reads were mapped and the read counts were calculated. Transcript expression and differentially expressed genes (DEGs) were determined using the EdgeR GLM approach (Robinson et al., 2010), with genes having FDRs < 0.01 classified as DEGs. Scaled expression values (fold change) were utilized for SOM-based clustering (Wehrens and Buydens, 2007). To assess the enrichment of Gene Ontology (GO) terms within the various sets of DEGs, GO enrichment analysis was conducted using Cytoscape with the BiNGO add-on (http://apps.cytoscape.org/apps/bingo) (Maere et al., 2005). The GO term finder evaluates the overrepresentation of GO categories using a hypergeometric test, with FDR correction for multiple testing (P ≤ 0.05). See Dataset S2 for specific gene sets. All RNA-seq data have been deposited to DDBJ with accession number PRJDB18465.

### Recombinant protein expression and purification

MPK6ΔNt (29-305) (Putarjunan et al., 2019), MPK6-CA_D218G_E222A_, MPK6-IA_K92MK93R,_ MKK5_DD_ and SCRM (1-494) were cloned into pGEX-4T-1 vector with an N-terminal GST tag and a thrombin cleavage sequence. For protein expression, the constructs were transformed into *E. coli* strain BL21. For each transformant, a single colony was selected and incubated in 10 ml LB liquid medium. The overnight-incubated *E. coli* suspensions were transferred to 1L LB medium and incubated at 37 °C for around 2 h until the OD_600_ reached 0.4–0.6. The isopropyl β-D-1-thiogalactopyranoside was added to the cultures and the strains were incubated further 6 h. GST-fused proteins were purified using glutathione agarose resin. The soluble portion of the cell lysate was loaded onto a GST-Sepharose column. Non-specifically bound proteins were removed by washing the column with 20 mM Tris pH 8.0, 200 mM NaCl. The bound GST-fused protein was eluted with 10 mm glutathione and 20 mM Tris pH 8.0, 200 mM NaCl (pH 8.0). The GST-fused proteins were exchanged with phosphate-buffered saline buffer, and then the solution was treated with 50 μg of thrombin for 10–12 h at 16 ^ο^C. The GST portion of the protein was cleaved during thrombin digestion, and then the whole solution was reloaded onto the glutathione *S*-transferase column to obtain pure protein. The purified proteins were further purified by gel filtration on a Superdex-200 (GE) column using fast protein liquid chromatography and phosphate buffer (pH 7.2) as the eluent. The purity of the protein was checked by SDS-PAGE (should we provide supplementary image?).

### Isothermal Titration Calorimetry (ITC)

Binding of the small molecules kC9, kC9-3 to MPK6ΔNt (29-305), MPK6-CA_D218G_E222A_, MPK6-IA_K92MK93R,_ kC9-3 to MPK6ΔNt-ATP (29-305) and SCRM KiDoK peptide binding with MPK6ΔNt (29-305) in the presence and absence of kC9 was characterized at 25 °C using a Malvern PEAQ-ITC microcalorimeter. All protein samples were dialyzed overnight using PBS buffer containing 5% DMSO. Small molecules kC9 and kC9-3 was dissolved in PBS buffer containing 2% DMSO. Titrations were performed by injecting 1 × 0.5-μl and 17 × 2-μl aliquots of 1 mM kC9/ kC9-3 to 30 mM of protein in PBS buffer pH 7.4, containing 2% DMSO. All titrations were carried out at least twice. The raw data were corrected using buffer and protein controls and analyzed using the software supplied by the manufacturer.

### Biolayer interferometry (BLI)

The binding affinities of the kC9, kC9-3 to MPK6, MPK6-CA_D218G_E222A_, MPK6-IA_K92MK93R_ 6 and SCRM KiDok peptide binding with MPK6 in the presence and absence of kC9 were measured using the Octet Red96 system (ForteBio, Pall Life Sciences) following the manufacturer’s protocols. The SCRM KiDok and scrm-D KiDok peptides were custom synthesized and biotinylated (BioSynthesis). The optical probes coated with anti-GST or streptavidin were first loaded with 500 nM GST tagged proteins or biotinylated SCRM KiDok peptide before kinetic binding analyses. The experiment was performed in 96-well plates maintained at 25 °C. Each well was loaded with 200 μl reaction volume for the experiment. The binding buffer used in these experiments contained 1× PBS supplemented with 2 % DMSO, 0.02 % Tween20, 0.1 % BSA. The concentrations of the kC9/kC9-3 as the analyte in the binding buffer were 100 mM, 50 mM, 25 mM, 12.5 mM, 6.25 mM, 3.12 mM, 1.56 mM and the concentrations of the MPK6/MPK6:kC9 (1:1.5) as the analyte in the binding buffer were 150 nM, 75 nM, 37.5 nM, 18.75 nM, 9.37 nM, and 4.68 nM. There was no binding of the analytes to the unloaded probes as shown by the control wells. Binding kinetics to all seven concentrations of the analytes were measured simultaneously using default parameters on the instrument. The data were analyzed using the Octet data analysis software. The association and dissociation curves were fit with the 1:1 homogeneous ligand model. The k*_obs_* (observed rate constant) values were used to calculate *K*_d_, with steady-state analysis of the direct binding.

### Docking Modeling

Molecular docking of ATP and kC9 (**2**) and its analogues (kC9-3 and kC9-8) to the MPK6 (PDB ID: 6DTL) (Putarjunan et al., 2019) were carried out using the high ambiguity-driven biomolecular docking (HADDOCK) approach (de Vries et al., 2007; Dominguez et al., 2003). Ambiguous interaction restraints (AIRs) were selected based on preliminary AUTODOCK model (Morris et al., 2009) and ATP interacting residues with ERK2 kinase (Zhang et al., 2012a). After an initial rigid body docking, the top 1000 structures that had the best intermolecular energies were then sequentially subjected to semiflexible simulated annealing and explicit solvent refinement. The pairwise “ligand interface RMSD matrix” over all structures was calculated, and the final structures were clustered using an RMSD cut-off value of 3.5 Å for both ligand and protein. The clusters were then prioritized using RMSD and the “HADDOCK score” (weighted sum of a combination of energy terms). The structures with best HADDOCK score were used for representation.

### Kinase-Glo Luminescent Kinase Assay

A luminescence assay was used to monitor the phosphotransferase activity of MPK6 kinase in presence and absence of kC9, kC9 analogues and known kinase inhibitor U0126. When activated, MPK6 phosphorylated a peptide substrate by converting ATP to ADP. The kinase reaction was stopped and unreacted ATP was removed using the ADP-Glo reagent. Kinase detection reagent was added to convert ADP back to ATP, which was then converted to a luminescent signal by the luciferase reaction (Promega, catalog No. V6072). Firstly, MPK6 protein alone or together with MKK5_DD_ were incubated in 30 μL of reaction buffer: 25 mM Tris-HCl (pH 7.5), 12 mM MgCl_2_, 1 mM DTT, and 50 μM ATP. The mixtures were kept at room temperature for 30 min. In the second step, 5 μL solution from each of the first reactions was incubated with SCRM/scrm-D in 25 μL of reaction buffer: 25 mM Tris-HCl (pH 7.5), 12 mM MgCl_2_, 1 mM DTT, 50 μM ATP. The kinase reaction was conducted at room temperature (25−27 °C) for 30 minutes. DMSO was used as a negative control for small molecule inhibitors, and a system without ATP was used to measure background signals. Luminescence was measured using an Promega microplate reader. The data were analyzed using GraphPad Prism 10 software.

### *In vivo* MAPK phosphorylation activity assays

MAPK phosphorylation activity assays were conducted with slight modifications to the procedures described by Lee *et al*. (2015). Five-day-old Arabidopsis Col-0 seedlings, initially grown on 1/2MS media plates, were transferred to 1/2MS liquid media in 12-well cell culture plates and incubated overnight in a long-day growth room. The kC9 priming treatment was performed by adding kC9 stock to the liquid medium to achieve a final concentration of 50µM at 6 hours. Seedlings were briefly dried on Kimwipe paper, and immediately frozen in liquid nitrogen 1 hour after peptide treatment. The samples were ground in liquid nitrogen and homogenized in protein extraction buffer (100mM Tris pH7.5, 150mM NaCl, 10% glycerol, 20mM Sodium fluoride, 1.5mM Sodium orthovanadate, 2mM Sodium molybdate, 1mM PMSF, 1% Triton x-100, 1x Protease inhibitor cocktail, 1x Phosphatase inhibitor cocktail 2, 1x Phosphatase inhibitor cocktail 3, 10µM MG-132 (Sigma). Protein concentration was determined by using the Bradford assay (Bio-Rad) with standard BSA. Equally normalized protein samples were subjected to immunoblot analysis using Anti-Phospho-p44/42 MAPK (Erk1/2) (Thr202/Tyr204) (Cat # 9101S, Cell Signaling Technology) as the primary antibody and peroxidase-conjugated goat Anti-rabbit IgG (Cat # AB205718, Abcam) as the secondary antibody. The same immunoblot membrane were analysed using an Anti-actin antibody (Cat # AB230169, Abcam) as the internal loading control. Signal intensity of immunoblot was quantified using Fiji ImageJ.

